# The role of size in biostability of DNA tetrahedra

**DOI:** 10.1101/2023.03.06.531312

**Authors:** Javier Vilcapoma, Akul Patel, Arun Richard Chandrasekaran, Ken Halvorsen

## Abstract

The potential for using DNA nanostructures for drug delivery applications requires understanding and ideally tuning their biostability. Here we investigate how biological degradation varies with size of a DNA nanostructure. We designed DNA tetrahedra of three edge lengths ranging from 13 to 20 bp and analyzed nuclease resistance for two nucleases and biostability in fetal bovine serum. We found that DNase I had similar digestion rates across sizes but appeared to incompletely digest the smallest tetrahedron, while T5 exonuclease was notably slower to digest the largest tetrahedron. In fetal bovine serum, the 20 bp tetrahedron was degraded ~four times faster than the 13 bp. These results show that DNA nanostructure size can influence nuclease degradation, but suggest a complex relationship that is nuclease specific.

---

It is now well-established that DNA can be used as a construction material for different types of nanoscale structures.^1^ As the field of DNA nanotechnology matures, work is increasingly focused on biological applications of DNA nanostructures, mainly in biosensing and drug delivery.^2,3^ With these applications come questions on suitability of DNA nanostructures to withstand biological environments that often contain non-ideal buffer conditions as well as nucleases that can degrade DNA.^4,5^ To address the biostability of DNA nanostructures, various strategies have been tested including the modification of component DNA strands with chemical groups such as hexaethylene glycol or hexanediol groups,^6^ coating nanostructures with oligolysines,^7^ polycationic shells^8^ or block copolymers^9^ and using unnatural base pairs.^10^ The choice of DNA nanostructure has also been explored for the effect of DNA nanostructure shape on cell entry and the cellular pathways by which DNA nanostructures are taken up within cells^11^. In our own prior work, we found that inherent design and crossover frequency in DNA motifs can lead to sometime major changes in biostability.^12^ We previously suggested that frequent crossovers were likely providing physical obstructions for the binding and activity of nucleases. From this, we hypothesized that a similar enhancement of biostability could be brought about by reduced size of DNA nanostructures, which would also shrink accessible regions of double stranded DNA. Here, we designed and executed a study to test our hypothesis, and measure the effect of DNA nanostructure size on biostability. We designed, assembled, and purified three sizes of a DNA tetrahedron, and measured degradation when subjected to two different nucleases and fetal bovine serum (FBS).

DNA tetrahedra are perhaps the most often-used structures in DNA nanotechnology, especially in biosensing and drug delivery applications.^13–15^ The tetrahedra we used here is constructed from four synthetic DNA strands, each of which hybridize to defined regions of the other three strands (**Figure 1a**). Based on previous studies,^16^ we designed different sizes of the tetrahedron with edge lengths of 13 bp, 17 bp and 20 bp using components strands that are each 41 nucleotides, 55 nucleotides or 63 nucleotides respectively (**Figure 1b**). We assembled the DNA tetrahedron by mixing equimolar ratios of the component strands in Tris-Acetate-EDTA (TAE) buffer containing 12.5 mM Mg^2+^ and heating the mixture to 90 °C and quickly cooling it to 4 °C. We confirmed formation of the tetrahedra using non-denaturing polyacrylamide gel electrophoresis (PAGE) by comparing the structure to control lanes containing subsets of components strands (**Figure 1c** and **Figure S1**). We then purified the assembled DNA tetrahedra of all three sizes using a gel-based method^17^ to eliminate higher order aggregates and single strands that were not incorporated into the tetrahedron.

**Figure 1.**
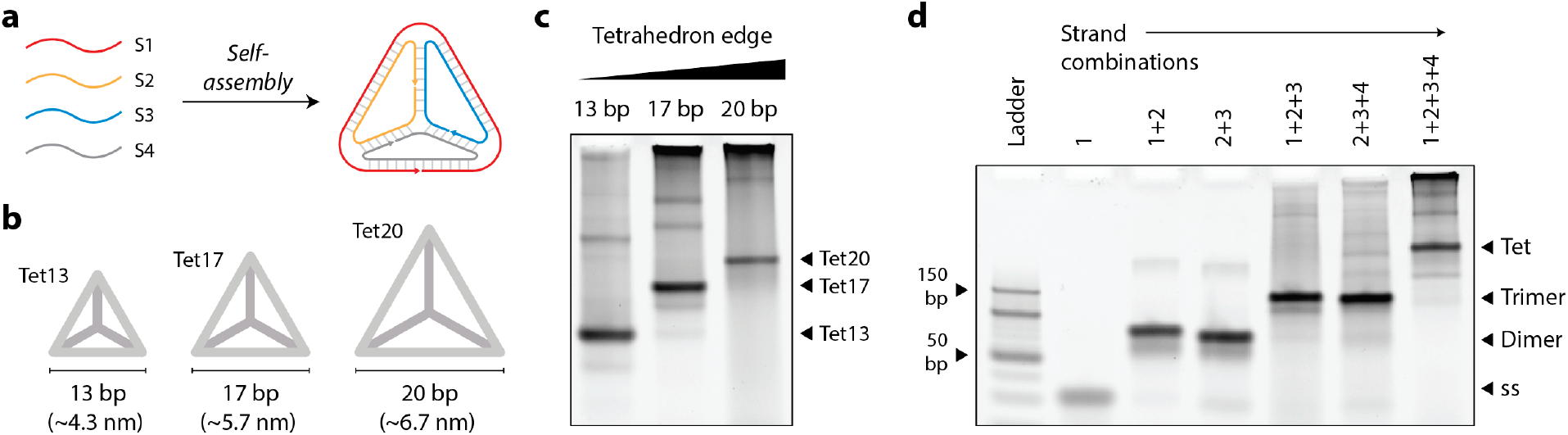
Assembly of DNA tetrahedra. (a) Self-assembly of four component strands into a DNA tetrahedron. (b) Tetrahedra with different edge lengths (13, 17 and 20 bp) constructed from pre-designed component strands. Duplex edges are simplified for clarity. (c) 10% non-denaturing PAGE showing the different sized tetrahedra as the major product in each lane. (d) Characterizing assembly of 17 bp DNA tetrahedra using 12% non-denaturing PAGE.

After validating formation of the DNA tetrahedra, we first tested nuclease resistance of these structures against DNase I. Since the optimal reaction temperature for DNase I is 37 °C, we first confirmed the thermal stability of the DNA tetrahedra at this elevated temperature (**Figure S2**). For nuclease resistance analysis, we incubated the assembled DNA tetrahedra with nucleases and analyzed degradation profiles on non-denaturing PAGE, a method we established before for other DNA nanostructures.^18^ We analyzed the kinetics of digestion by incubating the DNA tetrahedra with 0.01 U/μl DNase I and quantifying the band corresponding to the tetrahedron at different time points (**Figure 2a** and **Figure S3**). We had expected to see the smaller DNA tetrahedra to be more stable, assuming that the shorter edges occlude binding of the DNase I enzyme. However, the data seemed to indicate similar digestion rates with DNase I although the digestion seemed incomplete for the smallest size, perhaps indicating that some tetrahedra in the population do inhibit digestion. We performed a similar nuclease resistance experiment with another enzyme, T5 exonuclease (0.063 U/μl) (**Figure 2b** and **Figure S4**). Here we (surprisingly) observed the two smaller tetrahedra being degraded at similar rates while the largest tetrahedron degraded around 4-fold slower.

**Figure 2.**
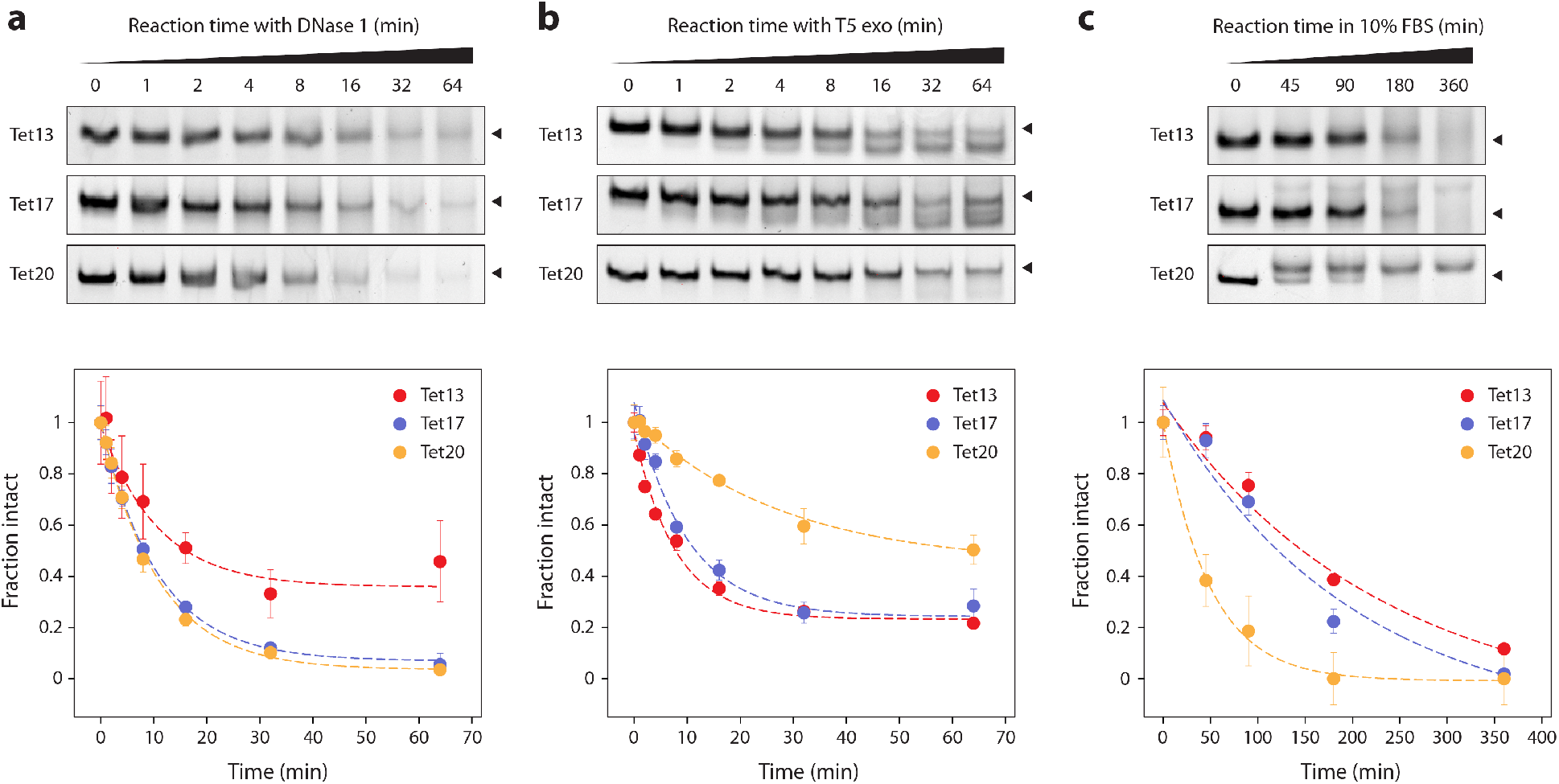
Biostability of DNA tetrahedra. Gel-based nuclease degradation analysis of tetrahedra with 13 bp, 17 bp and 20 bp edges in (a) 0.01 U/μl DNase I, (b) 0.063 U/μl T5 exonuclease and (c) 10% FBS. Bottom row shows quantified results from the gels in top row. Values represent mean and error propagated from standard deviations of experiments done in triplicates.

To provide a more realistic biological fluid, we incubated the DNA tetrahedra in 10% FBS at 37 °C for up to 6 hours (**Figure 2c** and **Figure S5**). This condition most closely matched our original expectation, showing that the larger tetrahedron degraded quickest and the smaller ones were more resistant to degradation.

Our study provides insight into the biostability of different sizes of the same nanostructure. While there is still a lack of consensus on which biostability enhancement strategies are better for different type of applications,^4^ obtaining data on the underlying biostability of DNA nanostructures will play an important role in tailoring these structures for various applications. In our study of the size correlation of DNA nanostructures to their biostability, our hypothesis seemed to be only partly true. Size dependence of nuclease degradation appears to be nuclease specific, and can strangely show opposite trends in different conditions. These effects could partly result from the different ways nucleases bind to their targets, and also potentially from where the terminal ends of strands reside. The structure of DNase I with duplex DNA shows that the enzyme requires 6-8 base pairs to bind to, and also typically causes a kink in the structure.^19^ For T5 exonuclease, while the nuclease shows exonuclease activity for double stranded DNA, ssDNA regions have been shown to thread through enzyme pockets.^20^ Thus there could also be differences in nuclease activity once the structure is acted upon by the enzymes.

Our hypothesis was too simplistic, and this work shows that biostability versus size cannot be easily generalized for different nucleases and solution conditions. Testing other nanostructures in the future will also provide data on whether these trends hold for other nanostructures, and also if there are differences between hollow nanostructures (similar to the tetrahedra studied here) and solid nanostructures. Our results can also inform on strategies to enhance biostability. In previous studies on DNA tetrahedra, activity of a restriction enzyme varied only when the nicks in the structure was ligated, and not in an unligated tetrahedron as the one used here.^21^ Further studies on a ligated system might yield more insights into enzyme action and in preventing or reducing degradation by nucleases when used in physiological conditions.

## Acknowledgments

Research reported in this publication was supported by the NIH through NIGMS under award R35GM124720 to K.H. and NIA under award R03AG076599 to A.R.C.

## Author contributions

JV performed experiments and analyzed data. AP performed experiments. ARC supervised the project, designed and performed experiments, analyzed and visualized data and wrote the manuscript. KH supervised the project, designed experiments, analyzed data and edited the manuscript.

