## Supporting Information for "The role of size in biostability of DNA tetrahedra"

### MATERIALS AND METHODS

#### Oligonucleotides

DNA strands were purchased from Integrated DNA Technologies (IDT). Full sequences (5'-3') are listed below.

Strands for tetrahedron with 13 bp edge (each strand is 41 nt):

```
TET13A: ACACTACGTCAGAACAGCTTGCATCACTGGTCACCAGAGTA
TET13B: ACGAGCGAGTTGATGTGATGCAAGCTGAATGCGAGGGTCCT
TET13C: TCAACTCGCTCGTAACTACACTGTGCAATACTCTGGTGACC
TET13D: TCTGACGTAGTGTATGCACAGTGTAGTAAGGACCCTCGCAT
```

Strands for tetrahedron with 17 bp edge (each strand is 55 nt):

```
TET17A: ACATTCCTAAGTCTGAAACATTACAGCTTGCTACACGAGAAGAGCCGCCATAGTA
TET17B: TATCACCAGGCAGTTGACAGTGTAGCAAGCTGTAATAGATGCGAGGGTCCAATAC
TET17C: TCAACTGCCTGGTGATAAAACGACACTACGTGGGAATCTACTATGGCGGCTCTTC
TET17D: TTCAGACTTAGGAATGTGCTTCCCACGTAGTGTCGTTTGTATTGGACCCTCGCAT
```

Strands for tetrahedron with 20 bp edge (each strand is 63 nt):

```
TET20A: AGGCAGTTGAGACGAACATTCCTAAGTCTGAAATTTATCACCCGCCATAGTAGACGTATCACC
TET20B: CTTGCTACACGATTCAGACTTAGGAATGTTCGACATGCGAGGGTCCAATACCGACGATTACAG
TET20C: GGTGATAAAACGTGTAGCAAGCTGTAATCGACGGGAAGAGCATGCCCATCCACTACTATGGCG
TET20D: CCTCGCATGACTCAACTGCCTGGTGATACGAGGATGGGCATGCTCTTCCCGACGGTATTGGAC
```

#### Formation of tetrahedron

To form the DNA tetrahedron, DNA strands A, B, C and D for each edge length were mixed in equimolar ratios in Tris-Acetic-EDTA-$Mg^{2+}$ (TAE/$Mg^{2+}$) buffer (1× final), which contained 40 mM Tris base (pH 8.0), 20 mM acetic acid, 2 mM EDTA, and 12.5 mM magnesium acetate. The DNA solution was heated to 90 °C for 5 minutes and quenched in ice for 5 minutes.

**Nuclease and FBS assay**

The annealed tetrahedra were mixed with DNase I reaction buffer or NE Buffer 2 at 1× final concentration for DNase and T5 exonuclease respectively. To this mixture, 0.01 U/μl DNase I or 0.063 U/μl T5 exonuclease were added at specific time points and incubated at 37°C. Dilutions of DNase I and T5 enzymes (New England Biolabs) to different units was made in nuclease-free water. Fetal bovine serum (FBS) was purchased from Thermo Fisher (Gibco, Catalog number: A4736301). For testing in FBS, the tetrahedra was mixed with FBS to a final concentration of 10% and incubated at 37°C for different time periods.

**Native polyacrylamide gel electrophoresis (PAGE)**

Non-denaturing gels containing 8-15% polyacrylamide (29:1 acrylamide/bisacrylamide) were run at 4°C (100 V, constant voltage) in 1× TAE/$Mg^{2+}$ running buffer. After electrophoresis, the gels were stained with GelRed (Sigma) and imaged using Bio-Rad Gel Doc XR+. Gel bands were quantified by integrating the peak areas using ImageLab. In the cases where multiple bands were not fully separable, we quantified the whole region and scaled by the ratio of the peak heights to estimate individual peak areas. Individual replicates were quantified separately, and each was normalized to its highest intensity. From the normalized individual replicates, we calculated the mean and SD for each time point. For the zero time point with value 1 we conservatively assigned the highest SD from the experiment and for points that decayed to exactly zero we assigned the lowest SD from the experiment. Final data was fit to single exponential decay curves with free fitting parameters and instrumental error weighting.

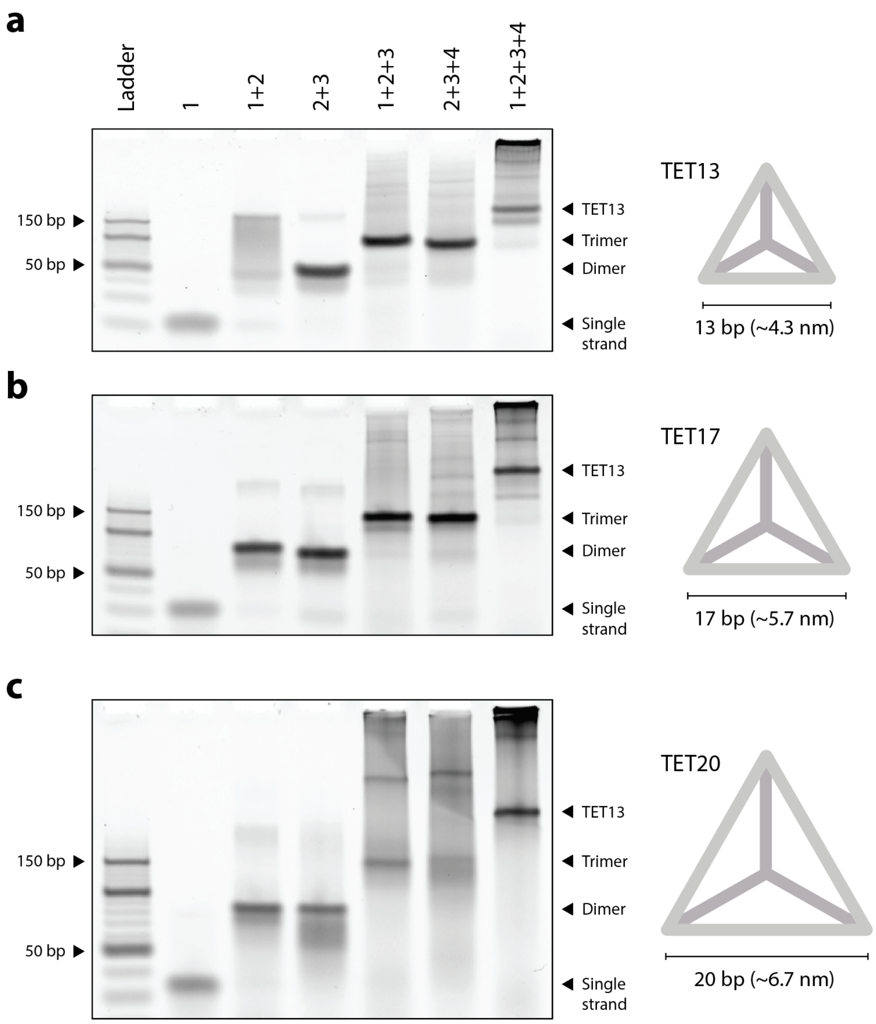


**Figure S1**. Characterization of DNA tetrahedra assembly using non-denaturing PAGE. (a) 13 bp, (b) 17 bp and (c) 20 bp.

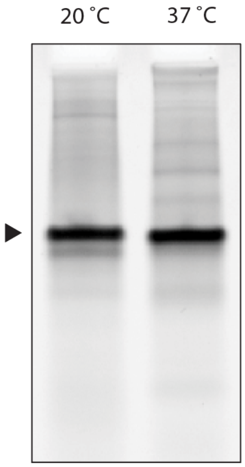


**Figure S2**. Stability of annealed tetrahedron at 37 °C for 2 hours.

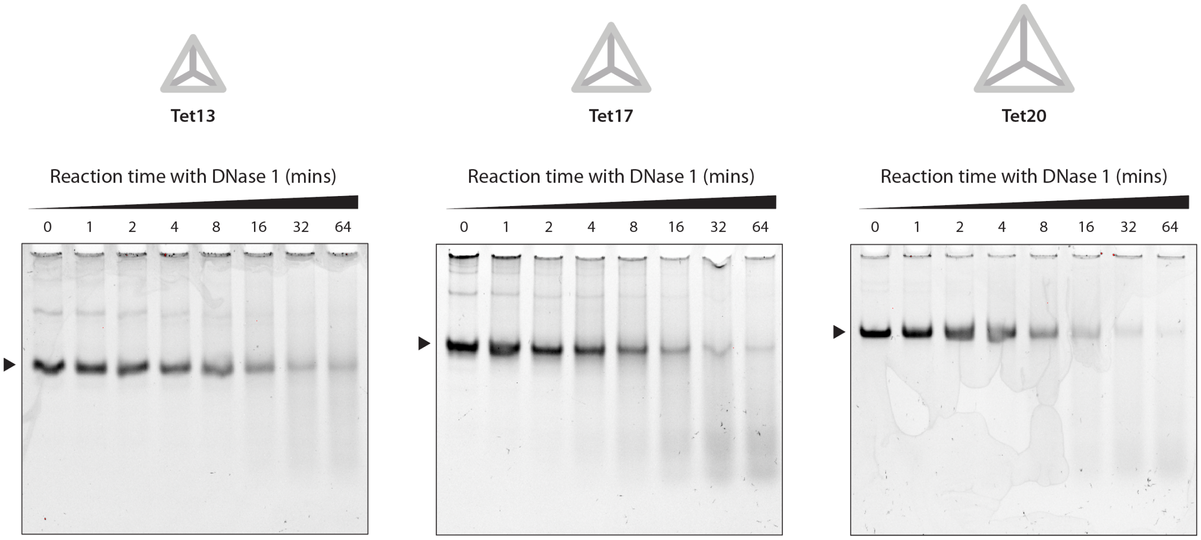


**Figure S3**. Time series of different sized tetrahedra in DNase I.

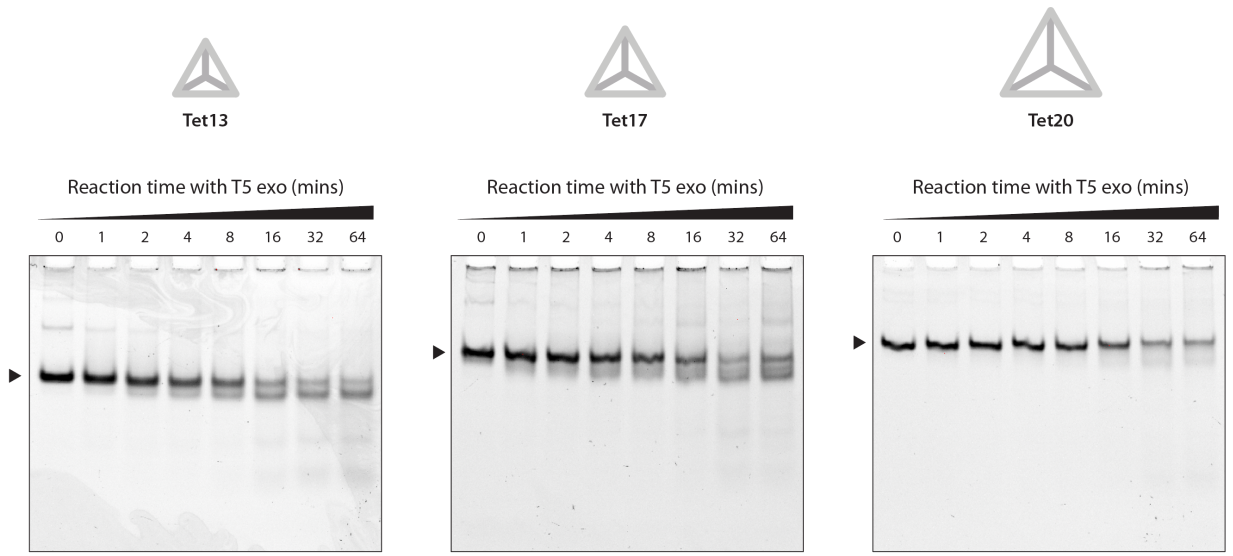


**Figure S4**. Time series of different sized tetrahedra in T5 exonuclease.

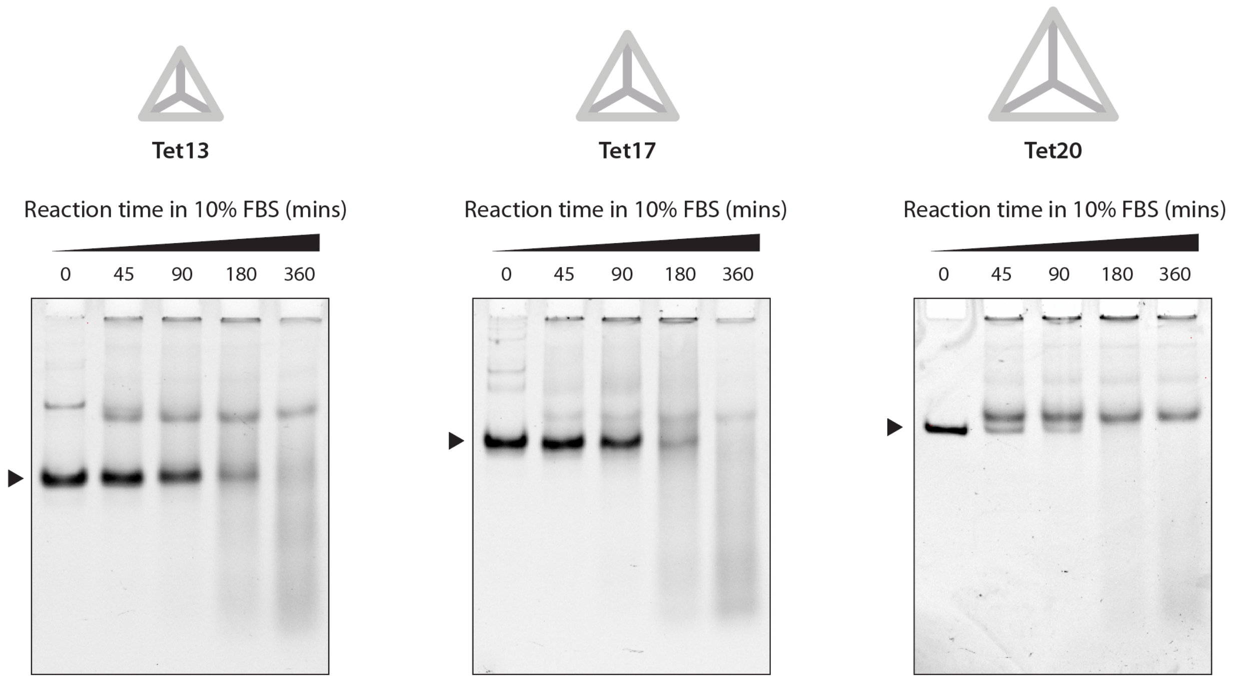


**Figure S5**. Time series of different sized tetrahedra in 10% FBS.
